# *Mycobacterium tuberculosis*-infected macrophages accumulate more nanoparticles

**DOI:** 10.1101/2025.05.15.654063

**Authors:** Amine Pochet, Tom Bourguignon, Axelle Grandé, Jesus Alfredo Godinez-Leon, Joan Fine, Priscille Brodin, Arnaud Machelart, Ruxandra Gref

**Affiliations:** Université de Lille, CNRS, INSERM, CHU de Lille, Institut Pasteur de Lille, U1019 - UMR 9017 - CIIL - Center for Infection and Immunity of Lille, F-59000 Lille, France; Université Paris-Saclay, CNRS, Institut des Sciences Moléculaires d’Orsay, 91405 Orsay, France

**Keywords:** PLGA, drug delivery systems, nanoparticles, tuberculosis, macrophages

## Abstract

In response to the escalating threat of infectious diseases and the widespread emergence of antimicrobial-resistant strains, novel therapeutic strategies are being actively developed. Nanoparticle-based delivery systems have emerged as promising tools to improve the targeted administration of antimicrobials directly to sites of infection. In the context of pulmonary infections, aerosolized nanocarriers can enhance local biodistribution while minimizing systemic exposure and associated side effects. In this study, we demonstrate that following pulmonary administration in *Mycobacterium tuberculosis*-infected mice, poly(lactic-co-glycolic acid) (PLGA) nanoparticles are broadly distributed throughout the lungs and preferentially interact with alveolar macrophages—the primary host cells of the pathogen. Remarkably, infected macrophages internalize significantly more nanoparticles than uninfected cells independently of the virulence and viability of the bacteria. Using transcriptomic analysis, we uncovered a list of 21 commonly modulated genes that may be responsible for this phenotype. Collectively, our findings reveal a striking feature of nanoparticles: their capacity to selectively accumulate in infected cells, offering a powerful mechanism to enhance antimicrobial delivery precisely where it is most needed.

## Introduction

Since their discovery, antimicrobials have revolutionized the treatment of infectious diseases and transformed modern medicine^1,2^. However, their extensive use—particularly in human health and agriculture^3,4^—has driven the rise of antimicrobial resistance (AMR), now recognized by the World Health Organization as one of the top ten global public health threats^5^. In response, new therapeutic strategies are being developed not only to combat existing resistance but also to prevent the emergence of new resistant strains^6,7^. One promising avenue is the use of drug delivery systems that enhance the targeted delivery of antibiotics to infection sites^8^. By allowing localized treatment via various administration routes (oral, parenteral, pulmonary, etc.), these systems can reduce drug dosages, treatment frequency, and off-target side effects, ultimately lowering the release of antimicrobials into the environment and helping preserve global health^9^.

Pulmonary administration is particularly attractive for treating respiratory infections due to the lungs’ extensive surface area and rich vascularization^10^. Nanocarrier-based strategies have already demonstrated improved bioavailability and efficacy of anti-infective agents in the lungs^11^. In the case of intracellular pathogens such as *Mycobacterium tuberculosis*, these systems can target alveolar macrophages—the primary host cells^12^—thereby enabling drug delivery in close proximity to the pathogen^13–15^. As such, nanocarrier-based antimicrobial delivery represents a powerful approach for localized treatment of infections and improving patient quality of life.

Among the available materials, nanoparticles based on poly(lactic-co-glycolic acid) (PLGA) stand out for their excellent biocompatibility, FDA approval, and widespread use in vaccinology and gene therapy^16–19^. In the field of infectious diseases, PLGA-based nanoparticles have been proven highly effective for encapsulating a wide range of anti-infective agents, with numerous well-documented preclinical studies supporting their use^20–24^. A key advantage of PLGA lies in its versatility: nanoparticles can be surface-functionalized to target specific cellular receptors or engineered to release their content in response to environmental cues, such as acidic pH, often found at infection sites^25–27^. In this study, we investigated the potential of PLGA nanoparticles for pulmonary drug delivery.

Using a murine model, we first demonstrated that PLGA nanoparticles distribute to distinct areas of the lung, where they primarily interact with alveolar macrophages. Importantly, in a *Mycobacterium tuberculosis in vitro* model of infection, we found that infected macrophages accumulate significantly more PLGA nanoparticles than their non-infected counterparts. Furthermore, this effect was consistent across different *M. tuberculosis* strains. Together, these findings reveal a key advantage of nanoparticles for targeted antimicrobial delivery: their intrinsic ability to preferentially accumulate in infected cells, offering a promising strategy to enhance the efficacy and precision of anti-infective therapies.

## Results and discussion

### PLGA nanoparticles accumulates in *M. tuberculosis*-infected alveolar macrophages

To investigate the behavior of PLGA nanoparticles in the lungs following pulmonary administration and their interactions with immune cell populations, nanoparticles were administered intranasally to C57BL/6 mice (50 µL containing 150 µg of PLGA) (**Fig. 1.a**). Three hours post-administration, histological analysis and flow cytometry were performed to evaluate these interactions. To enable direct fluorescence-based detection of the nanoparticles, rhodamine (Rho) was covalently conjugated to the terminal ends of the PLGA chains, thereby preventing dye leakage in biological media. The PLGA nanoparticles were round-shaped (**Fig. 1.b**) and had mean hydrodynamic diameters of 310 nm according to investigations.

**Figure 1:**
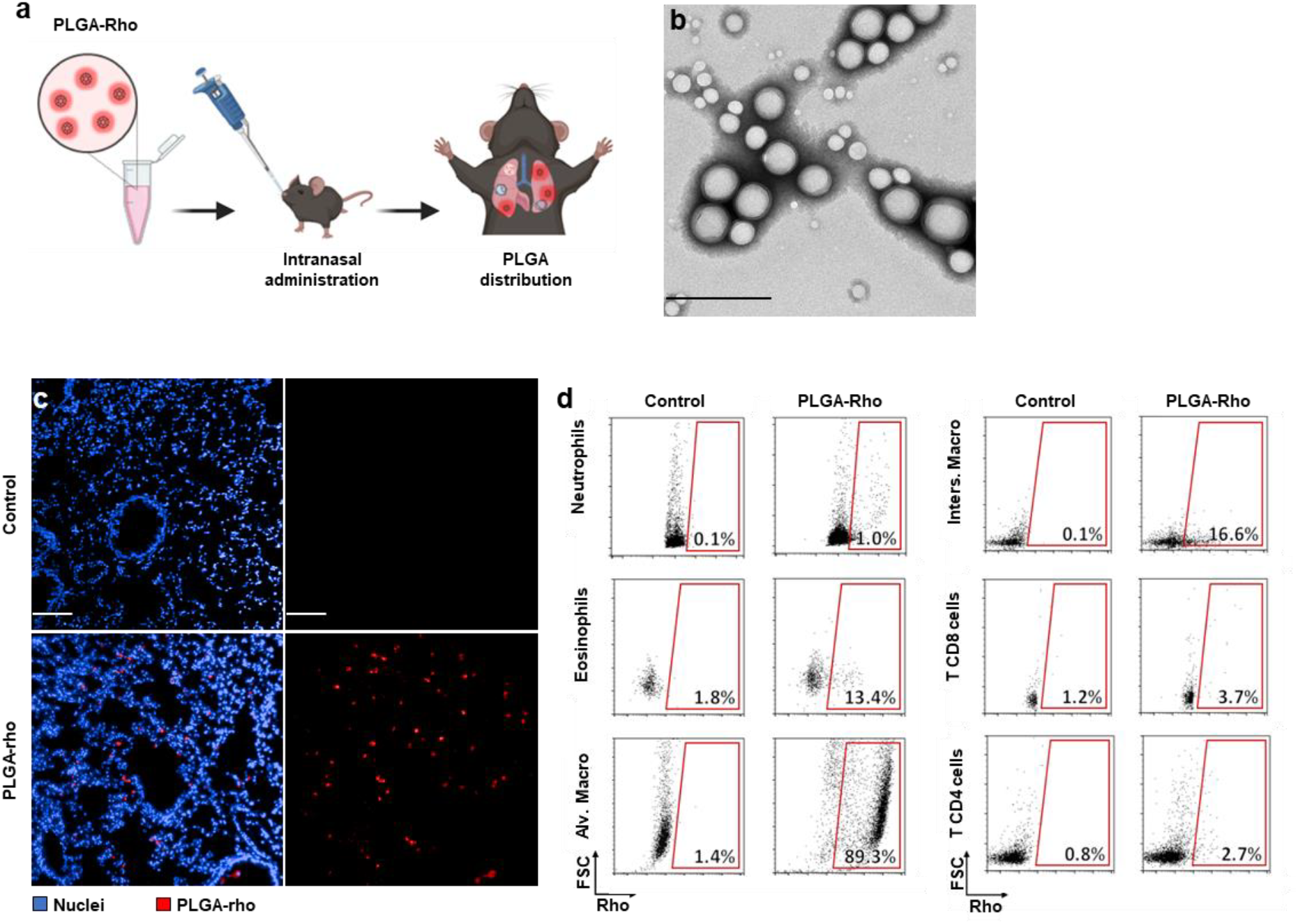
*M. tuberculosis* infected alveolar macrophages accumulate more PLGA than uninfected ones. **a**. Visual representation of the experimental procedure. Mice were administered NPs i.n. (150 µg in 50 μL) and euthanized 3 h post-administration to study NP distribution in lungs by histology and flow cytometry. **b**. Typical image obtained by TEM. Scale bar = 500 nm. **c**. Cryostat sections of lung tissues (10 µm) observed by fluorescence confocal microscopy, for untreated and treated mice (cell nuclei in blue, NPs in red). Scale bar = 50 µm. **d**. FSC (forward scatter) *versus* Rho plots of different cell populations extracted from untreated (control) and treated (PLGA-Rho) mice. N= 4 mice per group

The pulmonary distribution of the PLGA formulation after intranasal administration was investigated in mice using confocal microscopy. Lung cryosections (**Fig. 1.c**) revealed that PLGA nanoparticles disseminated deeply into the lung, reaching the alveolar spaces. Fluorescence imaging indicated heterogeneous accumulation patterns, suggesting either the formation of extracellular aggregates within alveoli or substantial cellular uptake. To clarify these observations, we analyzed the interaction of PLGA nanoparticles with various immune cell populations in the lungs using flow cytometry. The analysis demonstrated that multiple immune cell types interacted with the nanoparticles. Notably, 89.3 ± 2.32 % of alveolar macrophages (AMs) were PLGA-positive (**Fig. 1.d**), whereas other populations, including eosinophils and interstitial macrophages, showed limited association of 13.4% and 16.6%, respectively. Neutrophils, as well as CD4+ and CD8+ T cells, were largely negative for PLGA signal.

### M. tuberculosis infection enhances PLGA nanoparticle uptake by macrophages in vitro

To investigate nanoparticles interaction with infected macrophages, we established an *in vitro* model based on automated high-content imaging. In this context, high-content microscopy provides an ideal platform, allowing simultaneous visualization and quantification of multiple parameters. Acquired images are processed using dedicated software capable of distinguishing cellular components based on fluorescent signals. By developing tailored image analysis scripts and evaluating fluorescence colocalization, key parameters can be extracted, enabling the identification of distinct subpopulations, such as *M. tuberculosis*-infected cells^28^ (**Fig. 2.a**).

**Figure 2:**
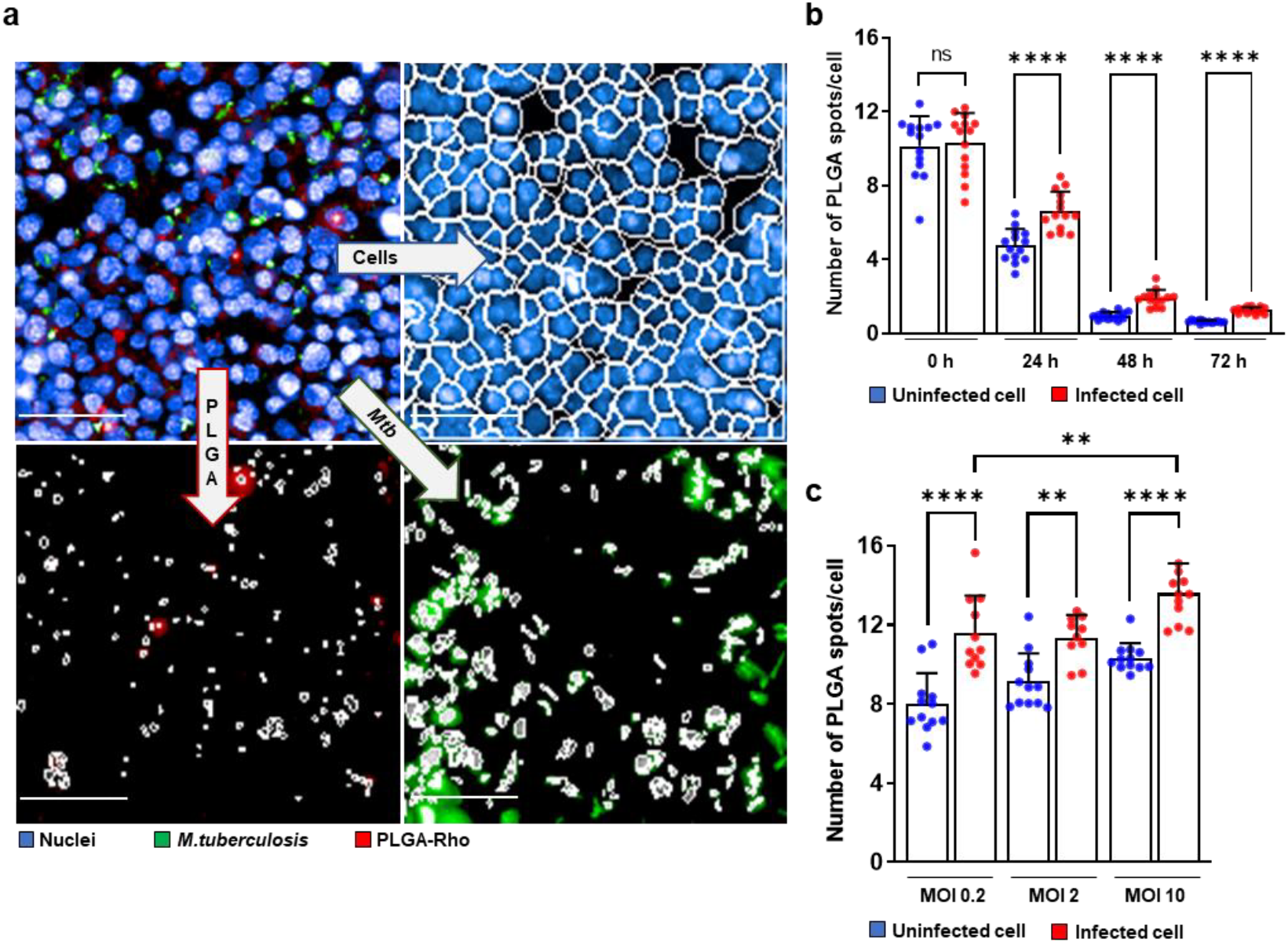
*In vitro M. tuberculosis* infected macrophages accumulate more PLGA than uninfected ones. **a**. Macrophages were infected by *M. tuberculosis (*MOI 2) in a 384-well plate. After 24 h of incubation, PLGA-Rho (20 µg/mL, 20 µL), diluted in cell culture medium, were then deposited into the wells. Finally, the plate was incubated for the desired period before being analyzed by high-content automated microscopy. The images are then processed using Columbus software, the input image (cell nuclei in blue, bacteria in green, NPs in red) allows distinct detection of PLGA-NPs, cells and bacteria based on their fluorescence. Scale bar = 100 µm **b**. Number of PLGA-rho spots per macrophage among infected and uninfected cells, depending on the period between their infection and their incubation with NPs (from 0 h to 72 h). **c**. Number of PLGA-Rho spots per macrophage among infected (red) and uninfected cells (blue), depending on MOI used. Data are presented as Mean ± SD and are representative of 3 different experiments. Ns: non-significant, ** P value < 0.01 and **** P value < 0.0001, as determined by Mann-Witney test.

To address this, PLGA nanoparticles were added at various time points from the moment of infection up to 72 hours post-infection. Interestingly, we observe that starting from 24 hours post-infection, the addition of nanoparticles leads to an increased accumulation in infected cells compared to non-infected ones. (**Fig. 2.b**). This suggests a delayed macrophage response to bacterial presence that enables increased nanoparticle uptake. Based on these findings, 24 hours post-infection was selected as the reference time point for subsequent experiments. To evaluate the influence of bacterial load on this process, macrophages were infected at increasing multiplicities of infection (MOI). Cells infected at higher MOIs exhibited significantly greater PLGA accumulation than those infected at lower MOIs (**Fig. 2.c**).

### Efficient accumulation of nanoparticles in *M. tuberculosis*-infected cells regardless of their size

The versatile physico-chemical properties of PLGA enables to produce nanoparticles with varying sizes, allowing us to investigate the influence of particle size on preferential uptake by infected cells. Five additional nanoparticle formulations were prepared and characterized by DLS (**Fig. 3.a**). The six resulting samples exhibited a wide range of mean hydrodynamic diameters, from 310 nm (± 100 nm) to 1035 nm (± 180 nm). Although the smallest batch appeared to be the least internalized, infected cells accumulated PLGA nanoparticles to a significantly greater extent than uninfected cells, regardless of particle size (**Fig. 3.b**).

**Figure 3:**
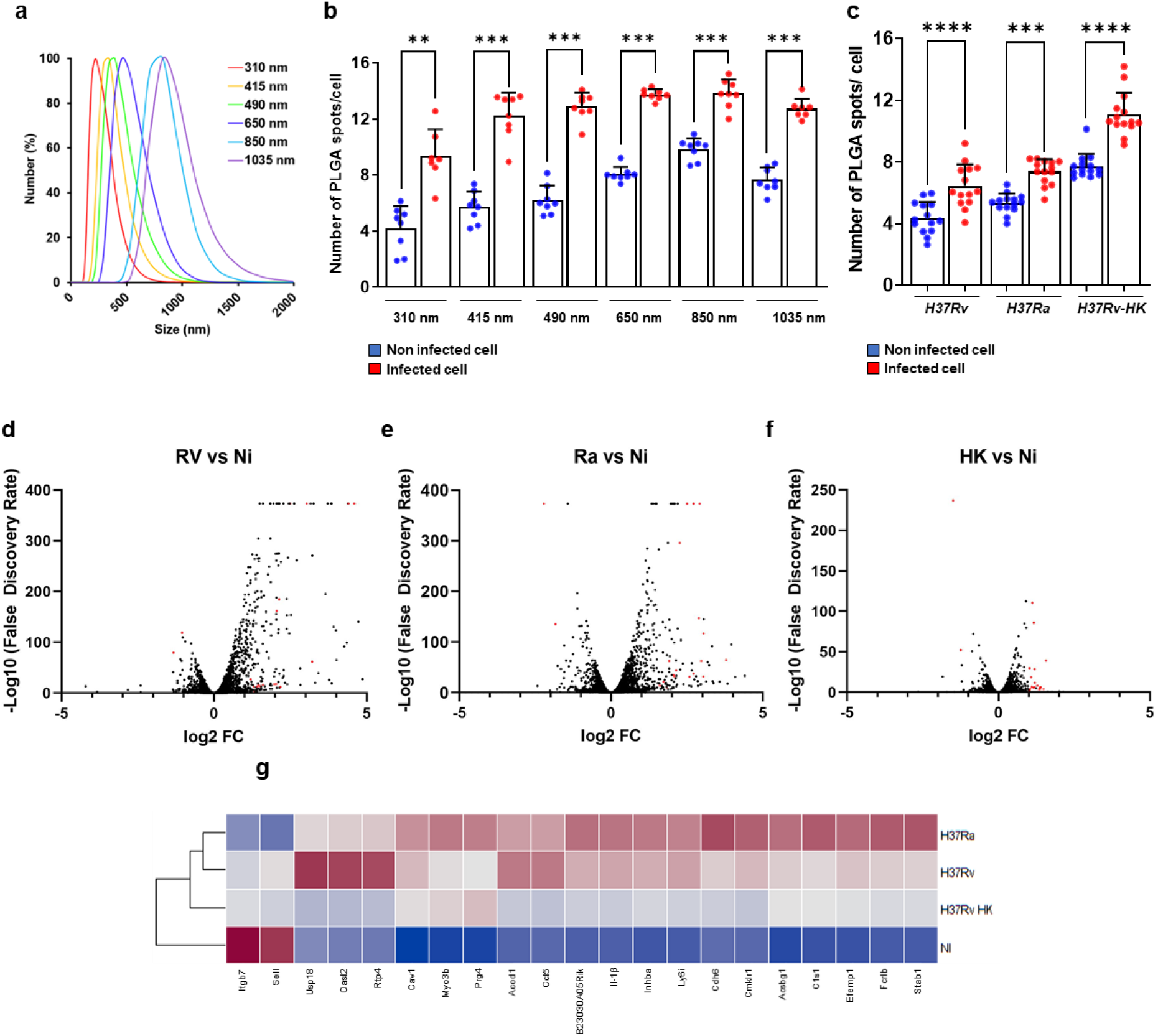
*M. tuberculosis*-infected cells accumulate more PLGA NPs regardless of the size and strain used. **a**. DLS size distribution profiles (number distribution) obtained for the six NP samples. The values are displayed as percentages to equalize the height of all six curves. **b**. Number of PLGA-rho spots among macrophages in infected (red) and uninfected cells (blue) depending on the size of the PLGA used. **c**. Number of PLGA-rho spots among macrophages in infected and uninfected cells depending on the strain of *M. tuberculosis* used. **d-g**.Total RNA was extracted from MPI cells after 24 h of infection with H37Ra, H37Rv, H37Rv-HK and sequenced using the illumina system. EdgeR analyses were then performed **d**. Volcano plots (Rv vs Ni) of MPI macrophages infected by H37Rv and control. **e**. Volcano plots (Ra vs Ni) of MPI macrophages infected by H37Ra and control. **f**. Volcano plots (HK vs Ni) of MPI macrophages exposed to H37Rv-HK and control. **g**. Differential expression of modulated genes in MPI cells infected with *M*.*tuberculosis* H37Rv, H37Ra, heat-inactivated H37Rv and uninfected cells. Each column represents the mean of 4 samples. The heat map was created using the Z-score of the normalized counts. Data are presented as Mean ± SD and are representative of 3 different experiments. Ns: non-significant, ** P value < 0.01, *** P value < 0.001 **** P value < 0.0001, as determined by Mann-Witney test

### Preferential nanoparticle accumulation occurs independently of bacterial virulence

*M. tuberculosis* produces several virulence factors, such as ESAT-6, which are known to modulate host intracellular pathways^29^. We hypothesized that these bacterial effectors could influence macrophage activity and thereby affect nanoparticle internalization. To test this, macrophages were infected with an attenuated strain of *M. tuberculosis* (H37Ra), known to secrete significantly fewer virulence factors^30^. In parallel, cells were also exposed to heat-killed *M. tuberculosis* (H37Rv-HK), which cannot actively produce or secrete any effectors. Following infection or stimulation, cells were incubated with PLGA nanoparticles under the same conditions as previously described. Surprisingly, in both conditions, infected or stimulated cells continued to exhibit preferential accumulation of nanoparticles compared to uninfected controls (**Fig. 3.c**). These results suggest that the phenotype is not driven by bacterial virulence or active effector secretion, but rather by the presence of mycobacteria within the host cells.

### Comparative transcriptomic analysis reveals limited shared gene modulation across *M. tuberculosis*-infection models

Since the preferential accumulation is independent of bacterial virulence, we used this observation to further investigate the mechanisms underlying this phenomenon. We compared the transcriptomic profiles of macrophages infected with either a virulent strain, an avirulent strain, or heat-inactivated bacteria, alongside non-infected control macrophages. The goal was to identify genes that are commonly modulated across the three infection conditions. RNA-seq analysis revealed a broad spectrum of differentially expressed genes across the various conditions. However, only 21 genes, comprising 19 upregulated and 2 downregulated, were commonly modulated in all three infection models (**Fig. 3.d-g**). Some of these genes are involved in the inflammatory response, such as Il-1β and Acod1. In summary, this transcriptomic analysis identified few commonly modulated genes, making it challenging to pinpoint a key pathway underlying this remarkable phenotype.

## Conclusion

In this study, we investigated the interaction between PLGA nanoparticles and *M. tuberculosis-*infected host cells. Using an infection model with *Mycobacterium tuberculosis*, we demonstrated that these nanoparticles exhibit a remarkable ability to preferentially accumulate in infected cells compared to non-infected ones. This property holds strong potential for improving antibiotic delivery directly to the infectious niche.

High-content imaging allowed us to gain deeper insights into the mechanisms and scope of this phenomenon. We observed that preferential accumulation is not limited to the virulent strain of *M. tuberculosis*. These results open up promising avenues for the design of targeted nanomedicine strategies to treat tuberculosis. However, a better understanding of the underlying cellular mechanisms remains essential. Therefore, it may be of interest to determine if this effect is conserved across diverse intracellular pathogens, including bacteria, viruses, and parasites. Identifying additional pathogens that trigger preferential accumulation could enable the investigation of the 21 commonly modulated genes under these conditions and potentially uncover a shared mechanism.

Several hypotheses should also be explored, including infection-driven alterations in host cell metabolism, cytoskeletal remodelling, endosomal trafficking disturbances, and membrane stress responses. Unravelling these pathways will be critical to fully harness this phenomenon and guide the development of next-generation, host-directed therapies using nanocarriers.

A key remaining question is whether infected cells accumulate greater amounts of encapsulated antibiotics. If confirmed, this could support the possibility of lowering the required drug dose, shortening treatment duration, and reducing systemic side effects. Collectively, our findings highlight the strong potential of antibiotic-loaded nanocarriers to selectively deliver therapeutic agents into infected cells, preferentially targeting those with higher bacterial burdens.

## Methods

### Materials

The biodegradable PLGA (50/50) copolymer was purchased from Expansorb (Aramon, France). PLA acid-terminated (MW = 18–24 kDa), piperazine, trimethyl aluminum (AlMe_3_), 4-dimethylaminopyridine (DMAP), 1-ethyl-3-(3-dimethylaminopropyl) carbodiimide (EDC), dichloromethane (DCM), Rhodamin B, polyvinyl alcohol (PVA) (88% hydrolyzed), RBS™ 35 concentrate and formalin solution (10%) were purchased from Sigma-Aldrich (Saint-Quentin-Fallavier, France). Mouse RAW 264.7 macrophages were purchased from ATCC (TIB-71™). Dulbecco’s phosphate buffered saline (PBS), Roswell Park Memorial Institute (RPMI) medium, fetal bovine serum (FBS) and Versene (ethylenediaminetetraacetic acid (EDTA)) were purchased from Gibco (Illkirch, France). Difco Middlebrook 7H9 was purchased from Becton Dickson (Grenoble, France). High purity glycerol was purchased from Euromedex (Souffelweyersheim, France). Oleic acid, albumin, dextrose and catalase (OADC) growth supplement was purchased from Fisher Scientific (MA, USA). Tween 80, genistein, cytochalasin D, wortmannin, chlorpromazine hypochloride and collagenase I were purchased from Sigma-Aldrich (Saint Louis, MO, USA). Hoechst and fixable viability dye Aqua were purchased from Fisher Scientific (Dreieich, Germany). CellBrite® cytoplasmic membrane red dye was purchased from Biotium (San Francisco, CA, USA). Collagenase was purchased from Roche (Switzerland). Ethanol 96% was purchased from VWR Chemicals (Fontenay-sous-Bois, France). The following monoclonal antibodies: fluorescein (FITC)-coupled HL3 (anti-CD11c), FITC-coupled 145-2C11 (anti-CD3), APC-coupled RB6-8C5 (anti-GR1), phycoerythrine (PE)-coupled RM4-5 (anti-CD4), BV650-coupled 53-6.7 (anti-CD8a), PE-coupled E50-2440 (anti-Siglec-F), and APC-coupled BM8 (anti-F4/80), as well as 2.4G2 (Fc Block CD16 CD32) and PermWash 10X, were purchased from BD Biosciences (Franklin Lakes, NJ, USA). APC-eF780-coupled M1/70 (anti-CD11b) was purchased from eBioscience (Villebon-sur-Yvette, France). Tissue-Tek® O.C.T.™ compound was purchased from Sakura (Flemingweg, Netherlands). Fluoro-Gel medium was purchased from Electron Microscopy Sciences (Hatfield, PA, USA). Pure water used in all experiments was filtered (18.4 MΩ·cm) by a Milli-Q system (Millipore, Milford, MA, USA). Goat serum was purchased from eurobio scientific (Les Ulis, France). Mouse anti-Influenza A Nucleoprotein antibody was purchased from Monosan (Uden, Netherland).

### NP preparation

To prepared fluorescent NPs were prepared as previously reported^31^ . Briefly, Rho was grafted to PLA, at the terminal carboxyl chain end of the polymer by adapting a previously described method^31,32^. Into 17.5 mL of a 0.22 mg/mL solution of piperazine in DCM were added 11.3 mL of a 2 M solution of AlMe_3_ in toluene. The resulting mixture was refluxed with Rho (0.5 mg/mL) for 24 h at 50°C and under inert atmosphere. The obtained Rho-piperazine amide was reacted with PLA acid-terminated using DMAP as catalyst and EDC as coupling agent.

PLGA-Rho NPs were prepared by emulsification-solvent evaporation. 150 mg of PVA were dissolved in 30 mL of Milli-Q water in order to prepare an aqueous solution of PVA (0.5% w/v). To obtain an aqueous suspension of NPs of 15 mg/mL, 60 mg of PLGA were dissolved in 1.5 mL of DCM, and 0.2 mg of PLA-Rho were added under magnetic stirring. Afterwards, 4 mL of the PVA were added to the solution, and the mixture was vortexed for 20 s to form a coarse emulsion. A finer emulsion was further obtained by two successive sonication steps: a first one of 90 s (20% power), and a second one of 30 s (10% power), using a sonicator probe. The solvent (DCM) was finally evaporated overnight under magnetic stirring.

### NP characterization

DLS was employed to determine NP mean hydrodynamic diameter, using a Malvern Zetasizer (Nano ZS90, Malvern Panalytical, Worcestershire, UK), with an equilibration time of 120 s. The samples were diluted in Milli-Q water (for a concentration in the range of 100 µg/mL), and their diameters were reported as Z_average_ (nm) ± standard error (SE). All experiments were performed in triplicates.

TEM analyses were performed on a JEOL JEM-1400 microscope operating at 80 kV. Onto copper grids covered with a formvar film (400 mesh), 5 µL of a NP suspension (of a concentration in the range of 10 mg/mL) were deposited for 60 s before the excess liquid was blotted off using filter paper. The samples were then stained using 2% phosphotungstic acid for 30 s and dried before observation. Images were acquired using a post-column high-resolution (9 megapixels) high-speed camera (RIO9, Gatan) and processed with ImageJ.

### Mice and ethics statement

Six-week-old C57BL/6NJ were purchased from Janvier (Le Genest-Saint-Isle, France). All experimental procedures were approved by the institutional ethical committee “Comité d’Éthique en Expérimentation Animale (CEEA) 75, Nord-Pas-de-Calais” and the “Education, Research and Innovation Ministry” (APAFIS#1 327 0232–2017061411305485 v6, approved on 09/14/2018). All experiments were performed in accordance with relevant guidelines and regulations.

### Cell culture

Raw 264.7 macrophages were cultivated until passage 12 in RPMI-FBS. For subsequent assays, cells were rinsed with D-PBS and harvested with Versene. After centrifugation, were resuspended into culture medium to be used.

### Administration of PLGA

C57BL/6NJ mice were anesthetized with a cocktail of xylazine (9 mg/kg) and ketamine (36 mg/kg) in PBS before being inoculated i.n. with PLGA-Rho NPs (in a volume of 50 µL), while control mice received 50 µL of water (vehicle).

### Flow cytometry

3 h post-administration, infected and non-infected mice, were sacrificed and their lungs were harvested. As previously described^13^, the lungs were then cut in small pieces and incubated for 1 h at 37°C in RPMI medium containing collagenase D (400 U/mL) and DNAse I (100 µg/mL). After incubation, fresh medium was added to stop the reaction of collagenase and DNAse I, and lung pieces were dissociated by suction and discharge. They were then filtered using a 100 µm SmartStrainer (Miltenyi Biotec) and centrifuged (350 g for 5 min).

Pellets were suspended in FACS buffer (PBS, 0.2% BSA, 0.02% NaN_3_) and transferred in 96-well plates. After another centrifugation (750 g for 2 min), cells were suspended in 200 µL of 2.4G2 diluted 200 times in FACS buffer, and stored at 4°C for 20 min to prevent antibody binding on the Fc receptor. Then, they were centrifuged (750 g for 2 min) and suspended in FACS buffer three times. Several combinations of fluorescent mouse antibodies diluted 250 times in FACS buffer were then used to stain lung cells (**Table** 1). After 30 min of incubation at 4°C, cells were washed three times and suspended in paraformaldehyde (1%) and stored for 15 min at 4°C. They were then washed three times following the same protocol and suspended in PermWash 1X for 10 min at 4°C. Finally, cells were washed two times and suspended in FACS buffer. The samples were analyzed using a NovoCyte Quanteon Flow cytometer (Agilent Technologies).

**Table 1:** Phenotypic determination of pulmonary immune cells.

| Cell type | Phenotype |
| --- | --- |
| Neutrophils | CD11b<sup>+</sup> Ly6g<sup>+</sup> |
| Eosinophils | F4/80<sup>+</sup> SiglecF<sup>+</sup> CD11c<sup>-</sup> |
| Alveolar macrophages | F4/80<sup>+</sup> SiglecF<sup>+</sup> CD11<sup>+</sup> |
| Interstitial macrophages | F4/80<sup>+</sup> SiglecF<sup>-</sup> CD11c<sup>+</sup> |
| T CD8 cells | CD3<sup>+</sup> CD8<sup>+</sup> |
| T CD4 cells | CD3<sup>+</sup> CD4<sup>+</sup> |

### Sample preparation for histology

After mouse sacrifice, lungs were fixed for 3 h at RT in a formalin solution (10%), washed in PBS, and incubated overnight at 4°C in a 20% PBS-sucrose solution under a vacuum. Tissues were then embedded in the Tissue-Tek® O.C.T.™ compound, frozen in liquid nitrogen, and cryostat sections of 10 μm were prepared. For staining, tissue sections were rehydrated in PBS and incubated for 30 min in a PBS solution containing Hoechst. Slides were mounted in Fluoro-Gel medium, and labelled tissue sections were visualized using the IN Cell 6500.

### Sample preparation for confocal microscopy

Cover glasses were sterilized by being immersed in a 2% RBS (cleaning agent) aqueous solution at 50°C, and in 90% ethanol afterwards. They were finally dried under a microbiological safety cabinet and placed at the bottom of 6-well plates. Infected cells were deposited into the said plates in a volume of 1 mL. In each well, 8×10^5^ macrophages and 1.6×10^6^ bacteria were deposited (MOI 2). The plates were stored in an incubator for 24 h (37°C, 5% CO_2_), so that cells could adhere and proliferate on the surface of the cover glasses. Then, RPMI medium was removed and replaced with 1 mL of fresh medium containing NPs (200 µg/mL). After a defined period of incubation, RPMI medium was removed and stored at 4°C for NTA studies. It was immediately replaced in the wells by a 4% formaldehyde solution for 30 min, before the fixation solution itself was removed, and the samples were washed three times with PBS and stored at 4°C. Before observation, cell nuclei were stained with a 1 µM Hoechst solution (30 min of incubation at 37°C), and cell membranes were stained with a CellBrite® Red solution (diluted 200 times in PBS, 2 min of incubation at RT). After staining, cells were washed three times with PBS.

Analyses were performed on a Leica DMI 6000 CS inverted confocal microscope, on a CUDA station equipped with the Leica Application Suite X (LAS-X) 3.5.6 software. The samples were studied using a 405 nm diode (Leica, 50 mW) and an Argon white laser WLL E (80 MHz, from 470 nm to 670 nm), on sequential mode, and a 63X PLAN APO oil immersion + DIC (differential interference contrast) objective (NA: 1.4) (Leica). Four detectors, two photomultiplier tubes (PMTs) (Hamamatsu 63557) and two hybrid detectors (HyD), were employed.

### *M. tuberculosis* infection assay

A total of 2*10^4^ macrophages were seeded in 384-well plates. The plates were stored in an incubator overnight (37°C, 5% CO_2_), to allow cell adhesion. Recombinant strains of *M. tuberculosis* H37Rv, H37Ra and expressing an enhanced green (H37Rv-GFP, H37Ra-GFP) were cultured at 37°C for 2 weeks in Middlebrook 7H9 medium (Difco) supplemented with 10% oleic acid-albumin-dextrose-catalase (Difco), 0.2% glycerol (Euromedex), 0.05% Tween 80 (Sigma-Aldrich), and 50 µg/mL hygromycin (Sigma-Aldrich). For infection, bacteria were washed 2 times in D-PBS and resuspended in RPMI-FBS. Clumped mycobacteria were removed by centrifugation at 700 rpm for 2 min and homogenous supernatant was used for infection. Bacterial titer was determined by measuring the optical density (DO 1 = 1*10^8^ bacteria/mL) and were added to macrophages and epithelial cells for 3 hours, respectively at a MOI of 2 and 50. 3 hours post-infection, cells were washed 3 times and incubated overnight in fresh medium. The following day, NPs were diluted to reach a suitable concentration (20 µg/mL) and deposited into the wells of interest in a volume of 20 µL. After 24 h of incubation, cells were fixed and labeled with 1µM Hoechst diluted in 10 % neutral buffered formalin solution for 30 min. After fixation, cells were washed using D-PBS and taken out of the microbiological safety cabinet for image acquisition.

### Image-based analysis

Images were acquired using an automated fluorescent confocal microscope (In Cell analyzer 6000, GE) equipped with a 20X (NA 0.70) air lens or 60X (NA 1.2) for pathogens infection intracellular mycobacterial replication and internalization of PLGA, pβCD and microsphere. The confocal microscope was equipped with 405, 488, 561 and 642 nm excitation lasers. The emitted fluorescence was captured using a camera associated with a set of filters covering a detection wavelength ranging from 450 to 690 nm. Hoechst 33342-stained nuclei were detected using the 405 nm laser with a 450/50-nm emission filter. Green signals corresponding to H37Rv-GFP and H37Ra-GFP were recorded using 488 nm laser with 540/75-nm emission filters. Red signals corresponding to PLGA and pβCD was recorded using 561 nm laser with 600/40-nm emission filters.

### Internalization pathway inhibition

Stock solutions of genistein, cytochalasin D and wortmannin were prepared using DMSO. Uninfected cells were deposited into 384-well plates as described above. After 24 h of incubation, RPMI medium was then replaced with the different internalization inhibitors diluted in fresh medium: genistein (125 µM), cytochalasin D (50 µM), wortmannin (12.5 µM). Cells were pre-incubated with inhibitors for 1 h before NPs were deposited. After 3 h of incubation with NPs, cells were washed with RPMI medium and fixed for 15 min using a 1 µM Hoechst solution diluted in a formalin solution (10%). Image acquisition was then performed using the IN Cell 6500 and Columbus system, in order to quantify the percentage of PLGA-positive cells.

### RNA extraction

Raw 264.7 were grown at 1*10^6^ cells/well in 6-well plates and RNA was extracted using QIAzol lysis reagent and miRNeasy Mini Kit according to the manufacturer instructions (Qiagen). RNA concentration was determined using the GE SimpliNano device (GE Healthcare, UK). Remaining DNA in samples was digested using the DNase I (Thermo Scientific, USA) for 20 min at RT.

### RNA sequencing

RNA quality was analyzed by the measurement of the RNA integrity number (RIN) with a bioanalyzer RNA 6000 Nano assay prior to sequencing. mRNA library preparation was realized following manufacturer’s recommendations (Ultra 2 mRNA kit from NEB). Final samples pooled library prep were sequenced on Novaseq6000 ILLUMINA with S1-200 cartridge (2×1600Millions of 100 bases reads) in one run, corresponding to 2×30Millions of reads per sample after demultiplexing. Quality of raw data was evaluated with FastQC. Poor quality sequences and adapters were trimmed or removed with the fastp tool, using default parameters, to retain only good quality paired reads. Illumina DRAGEN bio-IT Plateform (v3.8.4) was used for mapping on mm10 reference genome and for quantification using gencode vM25 annotation gtf file. Library orientation, library composition and coverage along transcripts were checked with Picard tools. Subsequent analyses were conducted with R software. Differential expression analysis was performed with the DESeq2 (v1.26.0) bioconductor package. Multiple hypothesis adjusted p-values were calculated with the Benjamini-Hochberg procedure to control FDR with a threshold of significance at 0.05. The cut-off for absolute log2-ratio was set at 1.5.

## Statistical analysis

Mann-Whitney and unpaired t-tests were applied using the GraphPad Prism software. Comparisons of groups two-by-two were performed, and the results are displayed when required. Values of *p* < 0.05 were considered significant. Indicated symbols of *, **, *** and **** denote *p* < 0.05, *p* < 0.01, *p* < 0.001 and *p* < 0.0001, respectively.

## Authorship contribution

R.G., A.M., P.B., T.B. and A.P conceived the study. A.P., T.B., A.M., A.G. and J. F. performed the experiments. R.G., A.M., T.B., A.P., A.G., J.F., B.D. and P.B. analyzed the data. T.B., R.G., A.M. and A.P. wrote the manuscript. All the authors approved the manuscript.

## Acknowledgments

We gratefully acknowledge Alexandre Vandeputte, Xue Lie, Jonathan Chatagnon, Berenice Dremierre, and Nathalie Deboosere for technical assistance and helpful discussions. We thank Nicolas Vandenabeele and Robin Prath for BSL-3 animal facility assistance. We thank the platform ARIADNE-criblage (UMS2014-US41 PLBS) to provide access to the high content microscope. We would like to thank Olivier Molendi-Coste and the Flow Core Facility of BioImaging Center Lille (UMS 2014 -US 41 - PLBS, F-59000 Lille, France) for the expert technical assistance. We would like to acknowledge Simang Champramary and the Paris Brain Institute’s Data Analysis Core for the RNA sequencing analysis (https://dac.institutducerveau-icm.org/). We would like to thank the management office of the Center for Infection and Immunity of Lille.

## References

1. Dhole, S., Mahakalkar, C., Kshirsagar, S. & Bhargava, A. Antibiotic Prophylaxis in Surgery: Current Insights and Future Directions for Surgical Site Infection Prevention. (2023).

2. Nnadozie, U. U. et al. Antibiotic use among surgical inpatients at a tertiary health facility: a case for a standardized protocol for presumptive antimicrobial therapy in the developing world. Infect. Prev. Pract. 2, 100078 (2020).

3. Mann, A., Nehra, K., Rana, J. S. & Dahiya, T. Antibiotic resistance in agriculture: Perspectives on upcoming strategies to overcome upsurge in resistance. Curr. Res. Microb. Sci. 2, 100030 (2021).

4. Menz, B. D. et al. Surgical Antibiotic Prophylaxis in an Era of Antibiotic Resistance: Common Resistant Bacteria and Wider Considerations for Practice. Infect. Drug Resist. 14, 5235–5252 (2021).

5. EClinicalMedicine. Antimicrobial resistance: a top ten global public health threat. eClinicalMedicine 41, (2021).

6. Blanco, P., Sanz-García, F., Hernando-Amado, S., Martínez, J. L. & Alcalde-Rico, M. The development of efflux pump inhibitors to treat Gram-negative infections. Expert Opin. Drug Discov. 13, 919–931 (2018).

7. Caradec, T. et al. A Novel Natural Siderophore Antibiotic Conjugate Reveals a Chemical Approach to Macromolecule Coupling. ACS Cent. Sci. 9, 2138–2149 (2023).

8. Ferreira, M. et al. Liposomes as Antibiotic Delivery Systems: A Promising Nanotechnological Strategy against Antimicrobial Resistance. Mol. Basel Switz. 26, 2047 (2021).

9. Kirtane, A. R. et al. Nanotechnology approaches for global infectious diseases. Nat. Nanotechnol. 16, 369–384 (2021).

10. Patil, J. S. & Sarasija, S. Pulmonary drug delivery strategies: A concise, systematic review. Lung India Off. Organ Indian Chest Soc. 29, 44–49 (2012).

11. Song, J. M. Intranasal delivery of liposomal indole-3-carbinol improves its pulmonary bioavailability. Int. J. Pharm. (2014).

12. Ahmad, F. et al. Macrophage: A Cell With Many Faces and Functions in Tuberculosis. Front. Immunol. 13, 747799 (2022).

13. Machelart, A. et al. Intrinsic Antibacterial Activity of Nanoparticles Made of β-Cyclodextrins Potentiates Their Effect as Drug Nanocarriers against Tuberculosis. ACS Nano 13, 3992–4007 (2019).

14. Nair, A. et al. Advanced drug delivery and therapeutic strategies for tuberculosis treatment. J. Nanobiotechnology 21, 414 (2023).

15. Costa-Gouveia, J. et al. Combination therapy for tuberculosis treatment: pulmonary administration of ethionamide and booster co-loaded nanoparticles. Sci. Rep. 7, 5390 (2017).

16. Allahyari, M. & Mohit, E. Peptide/protein vaccine delivery system based on PLGA particles. Hum. Vaccines Immunother. 12, 806–828 (2016).

17. Cappellano, G., Comi, C., Chiocchetti, A. & Dianzani, U. Exploiting PLGA-Based Biocompatible Nanoparticles for Next-Generation Tolerogenic Vaccines against Autoimmune Disease. Int J Mol Sci (2019).

18. Liu, C. et al. Folic Acid/Peptides Modified PLGA-PEI-PEG Polymeric Vectors as Efficient Gene Delivery Vehicles: Synthesis, Characterization and Their Biological Performance. Mol. Biotechnol. 63, 63–79 (2021).

19. Tobío, M., Gref, R., Sánchez, A., Langer, R. & Alonso, M. J. Stealth PLA-PEG Nanoparticles as Protein Carriers for Nasal Administration. Pharm. Res. 15, 270–275 (1998).

20. Mayattu, K., Rajwade, J. & Ghormade, V. Development of erythromycin loaded PLGA nanoparticles for improved drug efficacy and sustained release against bacterial infections and biofilm formation. Microb. Pathog. 197, 107083 (2024).

21. Guevara, A., Armknecht, K., Kudary, C. & Nallathamby, P. PLGA Nanoparticles Formulations Loaded With Antibiotics Induce Sustained and Controlled Antibiotics Release for Prolonged Antibacterial Action Against MRSA, and Pseudomonas aeruginosa FRD1. Mil. Med. 189, 230–238 (2024).

22. Landa, G., Aguerri, L., Irusta, S., Mendoza, G. & Arruebo, M. PLGA nanoparticle-encapsulated lysostaphin for the treatment of Staphylococcus aureus infections. Int. J. Biol. Macromol. 271, 132563 (2024).

23. Bourguignon, T. et al. Pulmonary delivery of clofoctol-loaded nanoparticles inhibits SARS-CoV-2 replication and reduces pneumonia. Int. J. Pharm. 677, 125634 (2025).

24. Bourguignon, T., Godinez-Leon, J. A. & Gref, R. Nanosized Drug Delivery Systems to Fight Tuberculosis. (2023).

25. Luan, H. et al. Mannosamine-Engineered Nanoparticles for Precision Rifapentine Delivery to Macrophages: Advancing Targeted Therapy Against Mycobacterium Tuberculosis. Drug Des. Devel. Ther. 19, 2081–2102 (2025).

26. Peng, C. et al. Mannosamine-Modified Poly(lactic-co-glycolic acid)-Polyethylene Glycol Nanoparticles for the Targeted Delivery of Rifapentine and Isoniazid in Tuberculosis Therapy. Bioconjug. Chem. (2025) doi:10.1021/acs.bioconjchem.5c00062.

27. Khanal, S. et al. pH-Responsive PLGA Nanoparticle for Controlled Payload Delivery of Diclofenac Sodium. J. Funct. Biomater. 7, 21 (2016).

28. Deboosere, N. et al. High-Content Analysis Monitoring Intracellular Trafficking and Replication of Mycobacterium tuberculosis Inside Host Cells. Methods Mol. Biol. Clifton NJ 2314, 649–702 (2021).

29. Anes, E., Pires, D., Mandal, M. & Azevedo-Pereira, J. M. ESAT-6 a Major Virulence Factor of Mycobacterium tuberculosis. Biomolecules 13, 968 (2023).

30. Jena, L., Kashikar, S., Kumar, S. & Harinath, B. C. Comparative proteomic analysis of Mycobacterium tuberculosis strain H37Rv versus H37Ra. Int. J. Mycobacteriology 2, 220–226 (2013).

31. Pancani, E. et al. High-Resolution Label-Free Detection of Biocompatible Polymeric Nanoparticles in Cells. Part. Part. Syst. Charact. 35, 1700457 (2018).

32. Nguyen, T. & Francis, M. B. Practical Synthetic Route to Functionalized Rhodamine Dyes. Org. Lett. 5, 3245–3248 (2003).

